# An optimal transport-based low-dimensional visualization framework for high-parameter flow cytometry

**DOI:** 10.1101/2025.08.19.670604

**Authors:** Abida Sanjana Shemonti, Grzegorz B Gmyrek, Katrien L A Quintelier, Sofie Van Gassen, Yvan Saeys, Marcella Willemsen, Joachim G J V Aerts, Eva V E Madsen, J Paul Robinson, Alex Pothen, Bartek Rajwa

## Abstract

Conventional data visualization techniques in single-cell analysis (such as two-dimensional dot plots, SPADE, PCA, t-SNE, or UMAP) often fall short in providing an intuitive understanding of high-parameter flow cytometry data. These methods tend to oversimplify complex biological relationships, lack biologically meaningful interpretations, and offer no principled framework for downstream quantitative analysis. To address these limitations, we present a graph-based visualization framework grounded in optimal transport theory. In this framework, cell populations are defined by their marker-expression profiles, and inter-population similarity is quantified using an efficiently computable optimal transport formulation known as the Sinkhorn distance. Our approach produces biologically consistent two-dimensional graph layouts using a phenotype-aware Hamming distance. Structural differences between sample graphs are characterized through a customized graph-edit distance that captures changes in population size, marker expression, and relationships between populations. We demonstrate our methods on two flow cytometry datasets: one from a clinical trial of dendritic cell-based immunotherapy in malignant peritoneal mesothelioma, involving 14 patients sampled at three time points with 14-color panels, and another from FlowCAP-II, which involved 43 acute myeloid leukemia patient samples analyzed with 7-color panels. Our framework produces robust, quantitative visual summaries of cell populations and supports statistical analysis based on graph edit distances, thereby offering new insights into disease progression and treatment response. Ultimately, our method bridges the gap between flow cytometry data visualization and biological interpretation.

## 1 Introduction

Flow cytometry (FC) is a high-throughput, multiparameter optical detection technology used to analyze the physical, biochemical, and functional properties of cells in a fluid suspension. By measuring thousands of cells per second, FC yields detailed information on cell size, internal complexity, and the presence of surface or intracellular markers (typically proteins) that define cell identity, activation state, and lineage. Modern instruments can assess up to 50 markers simultaneously (Konecny et al., 2024), enabling comprehensive profiling of multiple biological phenotypes in a single experiment. However, conventional two-dimensional dot-plot representations (defined in marker space) are inadequate for capturing the full complexity and richness of these high-dimensional datasets.

To address this limitation, the FC research community employs a range of computational approaches that form the backbone of modern visualization pipelines. These can be broadly grouped into three categories: (a) *dimensionality reduction* techniques that include linear methods such as principal component analysis (PCA) and multi-dimensional scaling (MDS), and non-linear manifold learning approaches such as t-SNE (t-distributed Stochastic Neighbor Embedding) (Van der Maaten and Hinton, 2008), UMAP (Uniform Manifold Approximation and Projection) (McInnes et al., 2018), and PHATE (Potential of Heat diffusion for Affinity-based Transition Embedding) (Moon et al., 2019); (b) *clustering* methods paired with graph-based algorithms such as SPADE (Spanning Tree Progression of Density Normalized Events) (Qiu et al., 2011), FlowSOM (Van Gassen et al., 2015), PhenoGraph (Levine et al., 2015), and X-shift (Samusik et al., 2016); and (c) inter-population similarity measures with *optimal transport (OT) framework* (Hauchamps et al., 2025; Gachon et al., 2025; Orlova et al., 2016). Although dimensionality reduction and clustering methods are well-established, they often oversimplify data, produce mathematically valid but biologically uninformative groupings, generate visualizations where distances between populations lack clear biological or statistical meaning, and are computationally demanding (see Figure 1).

**Figure 1.**
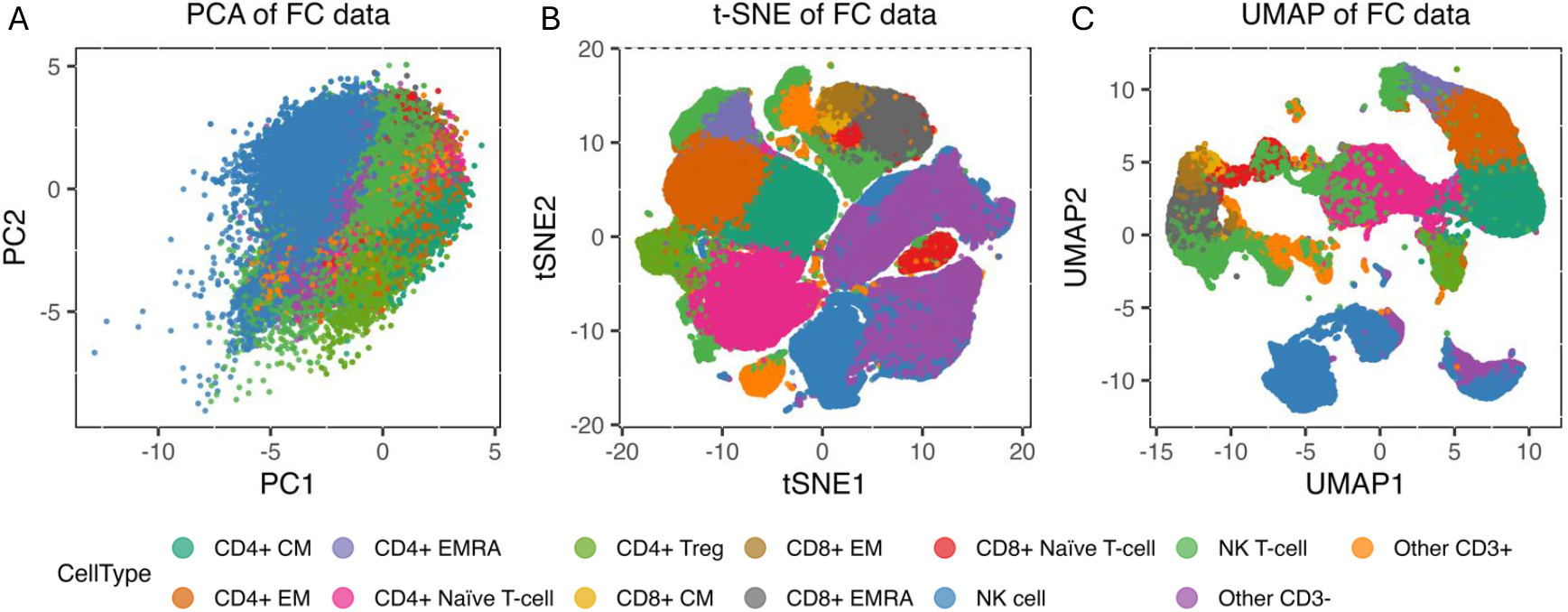
Three dimensionality reduction methods for flow cytometry data illustrating natural killer (NK) cells and T-cell populations. (A) PCA. (B) t-SNE. (C) UMAP. The plots are generated using the default set of parameters and the same flow cytometry sample (baseline sample of patient MCV005 in the malignant peritoneal mesothelioma dataset).

Previous studies have demonstrated the utility of high-dimensional FC combined with dimensionality reduction techniques to identify distinct immunotypes in patients (Mathew et al., 2020). In their work, Mathew et al. analyzed 200 immune features across healthy individuals, recovered and acutely ill COVID-19 patients, revealing three immunotypes with varying degrees of T- and B-cell activation that correlated with disease severity and clinical outcomes. These insights were primarily driven by visual exploration of immune marker distributions in UMAP and t-SNE space, highlighting the power of visualization in immunophenotyping. However, such projection-based methods can be limited in their ability to compare immune states across individuals or over time in a quantitative manner.

Despite its power to compare complete high-dimensional marker distributions, optimal transport (OT) remains underutilized in flow cytometry analysis, as only a handful of research groups have explored its potential. Orlova et al. (2016) were among the first to apply the Earth Mover’s Distance (EMD) to quantitatively compare biomarker-expression distributions across cell populations, revealing clinically relevant shifts. The optimalFlow framework (Del Barrio et al., 2020) extended this idea by clustering cytometry data and computing Wasserstein barycenters to create prototype templates, thereby improving supervised cell population identification despite biological and technical variability. More recently, CytOpT has used a regularized Wasserstein metric to estimate cell population proportions across samples while accounting for technical variation (Freulon et al., 2023). Similarly, Mukherjee et al. (2022) applied persistent homology to flow cytometry data from COVID-19 patients and healthy donors, leveraging selected marker combinations to construct persistence diagrams that capture topological features such as connected components and cycles. While this approach enables rigorous sample comparison via Wasserstein distances, the resulting persistent homology representations are neither easily interpretable in biological terms nor visually intuitive for clinical use. These pioneering studies highlight both the potential of OT for rigorous cytometry analysis and the need for broader adoption and improved interpretability in the field.

Although OT techniques have limited traction for analytical tasks in flow cytometry, their use in direct data visualization is even more limited. Notable efforts include CytoMDS, which combines the EMD with classical MDS to generate low-dimensional representations for visual quality control (Hauchamps et al., 2025), and the framework of Gachon et al. (Gachon et al., 2025), which integrates Wasserstein PCA and log-ratio PCA to produce biologically informed embeddings of high-parameter FC data, facilitating minimal residual disease detection and clustering in leukemia cohorts.

In this report, we introduce quantifiable visualization techniques for multiparameter FC data that are robust, mathematically rigorous, and biologically interpretable. Such methods are essential for conveying biological findings: they transform complex datasets into clear, actionable insights and facilitate the identification of patterns and relationships. Moreover, well-designed visualizations enhance communication and collaboration among researchers and clinicians by providing a shared framework for discussing results.

Our approach employs a regularized OT measure, known as the Sinkhorn distance (Cuturi, 2013). The principle is that each cell population’s phenotype is encoded by its marker-expression profile. Populations with similar marker abundances require less “work” to morph one distribution into another, yielding a smaller OT distance; conversely, dissimilar populations incur a higher transport cost. Accordingly, our OT framework defines transport costs directly in terms of the differences in marker abundances between populations. Classical OT computations can be prohibitively expensive for high-parameter data. The Sinkhorn distance overcomes this by introducing an entropic regularization term, which makes the optimization problem convex and enables the Sinkhorn–Knopp algorithm to converge rapidly to a near-optimal solution.

We encode inter-population Sinkhorn distances within graph-based visualization frameworks that condense the information contained in multiple two-dimensional dot plots and other conventional views into a single structure, which remains statistically informative for researchers and intuitive for non-technical users, such as clinicians. Our graph-based representation of FC samples also supports visual and quantitative comparisons of inter-sample differences via graph edit distances (GED). The key innovation of our methodology is the use of an efficiently computable OT measure to quantify similarity (or dissimilarity) between cell populations, enabling comprehensive and interpretable graph-based visualization tailored to monitor disease progression and treatment effects.

## 2 Materials and Methods

This section first details the flow cytometry datasets used to showcase our visual and quantitative analyses, then presents a clear, step-by-step description of the computational workflow.

### 2.1 Data acquisition and preprocessing

#### 2.1.1 Malignant Peritoneal Mesothelioma dataset

Flow cytometry data illustrating malignant peritoneal mesothelioma (MPM) is derived from a clinical trial of adjuvant dendritic cell–based immunotherapy (DCBI) conducted at Erasmus MC Cancer Institute in Rotterdam, Netherlands (Dietz et al., 2023). Fourteen patients were vaccinated at three time points, each two weeks apart, beginning 8–10 weeks after their cytoreductive surgery with hyperthermic intraperitoneal chemotherapy (CRS-HIPEC). Peripheral blood mononuclear cells (PBMCs) were collected from two cohorts (Group 1, N = 9; Group 2, N = 5) at baseline (pre-vaccination), two weeks post-first vaccine, and two weeks post-third vaccine. All samples were analyzed across six 14-color flow-cytometry panels and stored as FCS files.

Raw FCS files were processed in the R (v4.4.1) environment following the preprocessing pipeline of Dietz et al. Dietz et al. (2023). Briefly, we removed margin events and filtered the data using PeacoQC (Emmaneel et al., 2022), applied fluorescence signal unmixing (compensation), and rescaled marker abundances. Manual gating of Lymphocyte subpopulations was performed in FlowJo v10.10.0 (Becton, Dickinson and Company, 2023). Finally, data were harmonized using CytoNorm (Quintelier et al., 2025).

#### 2.1.2 Acute Myeloid Leukemia dataset

The Acute Myeloid Leukemia (AML) dataset originates from the FlowCAP-II initiative (Flow Cytometry: Critical Assessment of Population Identification Methods), which was established to benchmark and evaluate computational pipelines for distinguishing AML from healthy samples (Aghaeepour et al., 2013). This cohort comprises flow cytometry data from 359 individuals (316 healthy controls and 43 AML patients), acquired across eight 7-color panels and archived as FCS files. The raw data, already compensated and transformed by the repository, were further processed by scaling marker intensities and removing margin events before downstream analysis. The complete dataset is publicly accessible via FlowRepository: http://flowrepository.org/id/FR-FCM-ZZYA.

### 2.2 Cell population identification

Manual gating is the most traditional technique for identifying and quantifying specific cell populations in flow cytometry. It involves visually inspecting compensated fluorescence dot plots and delineating “gates” around populations of interest. Although intuitive and highly flexible, manual gating is labor-intensive and inherently subjective, which limits its scalability for high-throughput studies. Automated gating algorithms (including machine learning approaches) expand this framework and offer greater efficiency and reproducibility across large, complex datasets (Lux et al., 2018).

To manage the burden of manual annotation, researchers often perform gating or clustering in low-dimensional embeddings produced by manifold learning methods, such as t-SNE and UMAP. However, these projections can distort true inter-population relationships, reducing interpretability.

Unsupervised clustering techniques (such as DBSCAN (Ester et al., 1996), FlowSOM (Van Gassen et al., 2015), FlowGrid (Ye and Ho, 2019), and model-based approaches like LAMBDA (Abe et al., 2020)) provide another alternative to manual gates by grouping cells based on density or probabilistic models. Despite all these advances, many biologists still prefer manual gating for its direct control over population boundaries and its transparent, easily explained results.

Crucially, our visualization framework is agnostic to the upstream method of cell population assignment: it requires only that each cell be labeled, whether through manual gating, clustering, or other classifiers. While our examples assume discrete cell type labels, the approach naturally extends to probabilistic identities, in which each cell is represented by a vector of class-membership probabilities. We do not explore this extension here, as it falls beyond the current scope.

Our preprocessing employs a semi-automated, knowledge-driven pipeline for cell type assignment using a template of manually established gates. Beginning with a batch of normalized FCS samples and their associated cytometry panel, we first enumerate the candidate cell populations informed by marker panels and published phenotype definitions. From these descriptions, we compile a gating schema that specifies the marker–expression status of each population (e.g., positive vs. negative, high vs. low). We then apply this schema to one or a few representative samples, defining precise fluorescence-intensity thresholds (rectangular gate boundaries; see Figure 2). Because all samples are harmonized before processing, these thresholds transfer directly across the dataset.

**Figure 2.**
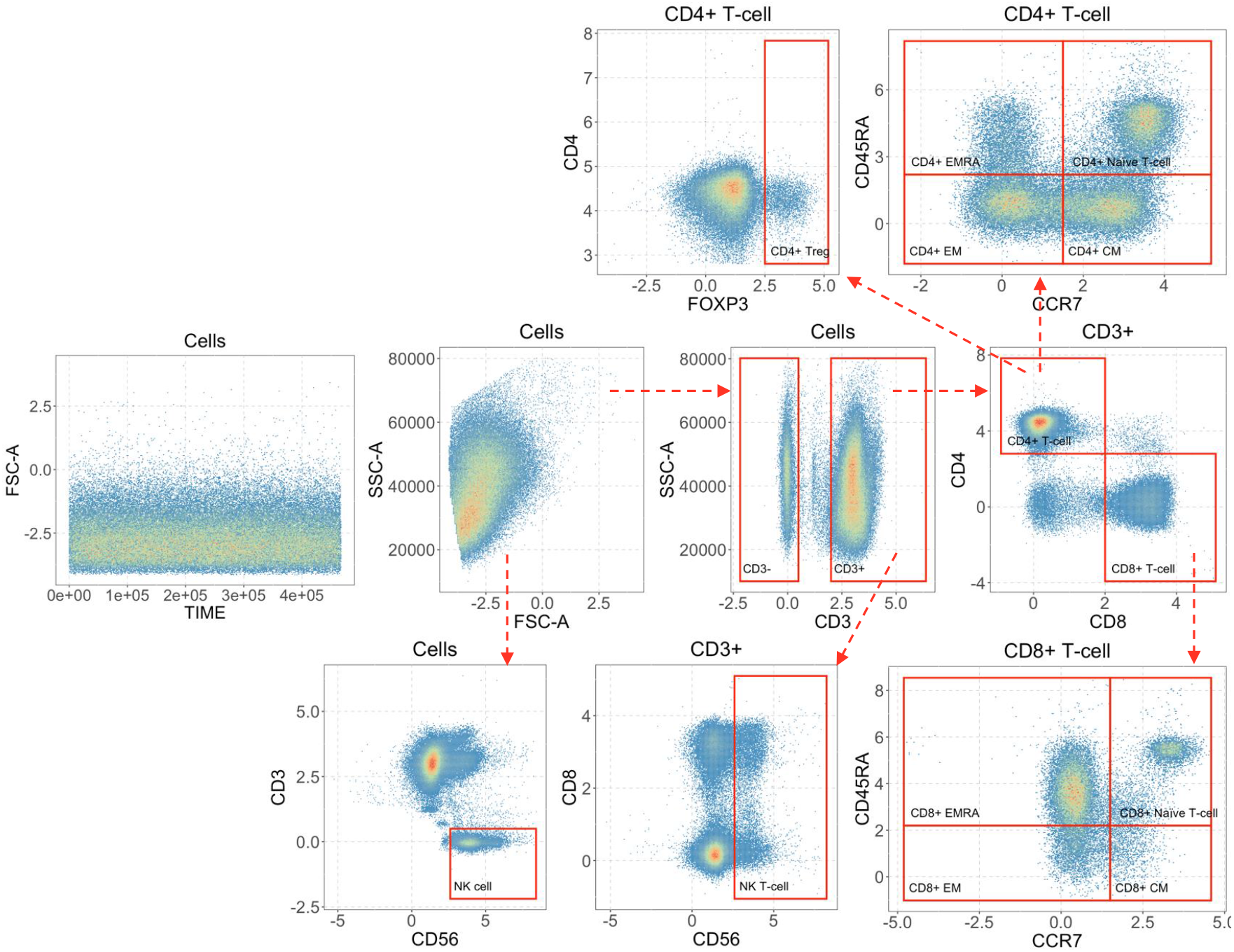
An illustration of the semi-automated gating process that identifies natural killer (NK) cell and T-cell populations in an MPM patient post-first vaccine (Patient ID: MCV005).

Next, we encode the gating thresholds as conditional statements in our processing workflow, automatically assigning each cell to a defined population based on its measured marker intensities. This approach is panel-specific, tunable, and transparent. It preserves expert knowledge rather than inferring clusters purely from data, yet it is far more efficient than fully manual gating. Importantly, as mentioned earlier, our downstream quantification and visualization steps are agnostic to the choice of cell-assignment method, provided that every cell carries a population label. Finally, these explicit phenotype definitions form the backbone of our visualization framework, as described in the following section.

### 2.3 Inter-population Sinkhorn distance computation

After identifying the cell populations in a cytometry sample, we calculate the Sinkhorn distances between each pair of populations. Full mathematical details on the optimal transport framework and the Sinkhorn distance computation are provided in the Appendix. Here, we offer a concise overview of the process.

In brief, consider two cell populations, *C*_1_ and *C*_2_, containing *n*_1_ and *n*_2_ cells, respectively. We assign each cell in *C*_1_ a mass of 1*/n*_1_ and each cell in *C*_2_ a mass of 1*/n*_2_, so that both distributions sum to one. Next, we construct the cost matrix

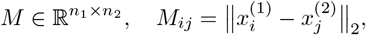

where 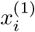 and 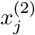 are the marker-expression vectors of cells *i* and *j* in *C*_1_ and *C*_2_, respectively.

We then solve the entropically regularized optimal transport problem (see the Appendix) using the Sinkhorn–Knopp algorithm to obtain the transport plan 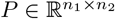. The Sinkhorn distance is given by

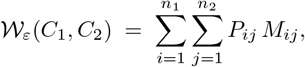

which quantifies the minimum “work” required to morph the distribution of *C*_1_ into that of *C*_2_. By normalizing each population’s total mass to one and leveraging entropic regularization for rapid convergence, this distance captures rich, high-dimensional differences in marker abundances, providing a rigorous and interpretable foundation for our visualization and comparison workflows.

### 2.4 Graph-based data visualization

Once we obtain the Sinkhorn distances between the cell populations, we visualize the cytometry samples using graphs. In this report, we adopt the standard mathematical definition of a graph *G* = (*V, E*), where *V* is the set of vertices (also called nodes) and *E* is the set of edges linking pairs of vertices. For example, in a social network graph, each node represents an individual, and each edge represents a friendship, enabling the analysis of connectivity, community structure, or influence. By the same convention, in our flow cytometry framework, each vertex corresponds to a distinct cell population.

Crucially, each edge in our graphs carries two attributes:

**1. Weight:** Visually encoded as edge thickness (and color), proportional to the inter-population Sinkhorn distance *W*_*ε*_(*C*_*i*_, *C*_*j*_), which quantifies distributional differences in marker expression between cell populations *C*_*i*_ and *C*_*j*_.

**2. Length:** Determined by the phenotype distance *d*_H_(*ϕ*_*i*_, *ϕ*_*j*_), computed via the Hamming distance (Hamming, 1950) between expert-defined phenotype strings (*ϕ*_*i*_ and *ϕ*_*j*_, for cell populations *C*_*i*_ and *C*_*j*_, respectively), so that populations with more similar marker-expression patterns lie closer together. The process of computing *d*_H_(*ϕ*_*i*_, *ϕ*_*j*_) is described below.

This dual encoding ensures that both the cost of “transporting” one distribution into another and the categorical phenotypic similarity are represented simultaneously and unambiguously.

For graph-based visualization, we leverage R’s *ggplot2* and *igraph* packages (Wickham, 2016; Csárdi et al., 2025). The *igraph* library supplies a high-performance suite for constructing, manipulating, and analyzing graphs, complete with numerous layout algorithms and network statistics. However, its built-in layout algorithms often rely on random initialization, leading to inconsistent representation across runs, which is problematic for comparative cytometry visualizations. To ensure both reproducibility and biological relevance, we instead compute domain knowledge-informed layouts, using Hamming distances to deterministically position vertices in a manner that reflects underlying cell phenotype relationships as understood by immunologists. By decoupling layout (Hamming) from edge-weight encoding (Sinkhorn), our approach guarantees both reproducibility and a clear, interpretable representation of phenotypic relationships.

**Phenotype-aware layout** Central to our visualization framework is the explicit encoding of expert-defined phenotype descriptors as fixed-length strings, one entry per marker in the cytometry panel. For example, suppose our panel comprises the markers

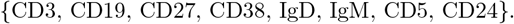

We represent the regulatory B-cell phenotype

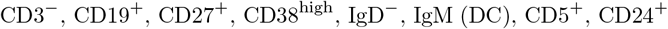

by the string [−, + +, ++, −, DC, +, +].

Here, “DC” (for “Don’t Care”) originates from Boolean algebra and digital-circuit design, where it denotes a variable whose value (0 or 1) does not affect the logical outcome. In our phenotype strings, “DC” indicates that the corresponding marker (IgM) is irrelevant for defining this population.

We then compute the *phenotype distance* between any two populations via the Hamming distance, but excluding positions labeled “DC” in either string. For instance, the strings

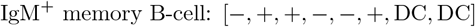

and

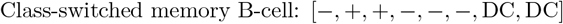

differ only at the IgM position (between “+” and “–”), yielding a phenotype distance of *d*_H_ = 1. Note that, the classical Hamming distance counts the number of differing positions between two strings. However, when computing phenotype distances between cell populations, we can customize these distances to reflect biological relevance. For instance, the difference between “+” and “++” can be set to 0.6, while the difference between “+” and “−” can be 1. In our experiments, we used uniform distances of 1 for all marker expression level pairs.

These pairwise phenotype distances form a symmetric matrix, denoted as *H*. To produce a reproducible, biologically informed two-dimensional layout, we apply the Kamada–Kawai algorithm (Kamada et al., 1989) to *H*. The Kamada-Kawai layout algorithm computes a force-directed layout by solving a nonlinear system of equations iteratively, yielding a fixed configuration of vertices (one per cell population) such that Euclidean separations approximate the phenotype distances. By anchoring each population at these deterministic coordinates, we eliminate layout variability and ensure that vertex positions carry direct interpretive meaning across all samples.

With vertex positions fixed, each cytometry sample is visualized as a weighted graph: vertex radii scale with the population’s fraction of total cells, and edges encode the inter-population Sinkhorn distances (Section 8) through thickness and color. This single graph representation unifies high-dimensional marker-expression profiles and population frequencies into a compact, interpretable summary, eliminating the need for multiple two-dimensional dot plots while preserving both biological and statistical insights.

### 2.5 Inter-sample GED computation

Up to this point, we have focused on methods for representing and visualizing individual flow cytometry samples. However, assessing the similarity or dissimilarity between pairs of samples can yield valuable insights, such as distinguishing healthy from diseased states, understanding disease mechanisms and progression, and evaluating treatment effects. Our graph-based representation enables inter-sample comparisons in a computationally efficient and biologically meaningful manner by leveraging the concept of graph edit distance (GED). GED quantifies the minimum cost required to transform one graph into another through a sequence of edit operations, such as adding, deleting, or substituting vertices and edges. As a well-established tool in pattern recognition, machine learning, and network analysis, GED provides a robust framework for comparing graph structures. However, because exact GED computation is NP-hard, we adapt the approximate GED implementation from Python’s *NetworkX* library (Hagberg et al., 2008), based on the method proposed by Abu-Aisheh et al. (2015), which applies depth-first search (DFS) with pruning over the space of partial edit paths.

In our approach, we enhance the biological relevance of GED by assigning custom costs to edit operations. Vertex edit costs reflect changes in the proportions of cell populations, while edge edit costs capture variations in inter-population Sinkhorn distances, which quantify differences in cell marker expression. This biologically informed cost model ensures that the computed GED between two graph representations accurately reflects the underlying biological variation between the corresponding flow cytometry (FC) samples. Moreover, prior knowledge of the cell populations reduces the complexity of the approximate GED computation. In such cases, GED can be computed efficiently by summing the substitution costs, resulting in a linear-time algorithm with respect to the number of vertices and edges.

This concludes the description of the computational components used in our method. The following sections present the resulting visual summaries and quantitative measures using the MPM and the AML dataset (described in Section 2.1).

## 3 Optimal transport-based visualization for MPM dataset

To illustrate our computational framework, we analyzed a subset of the MPM dataset using three cytometry panels optimized for T-cell profiling: *Co-inhibition, Co-stimulation*, and *Cytokine* panel.

- **Core cell-type markers (all panels):** CD56, CD3, CD4, FoxP3, CD8, CD45RA, CCR7, and Live/Dead viability channel.
- **Panel-specific cell-state markers:**
  – *Co-inhibition:* LAG3, PD-1, TIM-3, CD39, KI-67, CTLA-4
  – *Co-stimulation:* CD28, CD137, PD-1, HLA-DR, ICOS, KI-67
  – *Cytokine:* PD-1, TBET, IL-10, TNF-*α*, IL-2, IFN-*γ*

Using the shared core markers, our semi-automated gating procedure identified thirteen cell populations:

- Natural killer (NK) cells
- NK T-cells
- CD4^+^ regulatory T-cells (Tregs)
- CD4^+^ effector memory (EM) T-cells
- CD4^+^ terminally differentiated effector memory (EMRA) T-cells
- CD4^+^ central memory (CM) T-cells
- CD4^+^ naïve T-cells
- CD8^+^ EM T-cells
- CD8^+^ EMRA T-cells
- CD8^+^ CM T-cells
- CD8^+^ naïve T-cells
- Uncategorized CD3^+^ cells
- Uncategorized CD3^−^ cells

Please refer to Section 2.2 and Figure 2 for additional details.

Figure 3 illustrates four phenotype-aware two-dimensional layouts derived from the marker-expression profiles of our identified cell populations. To generate these, we first computed the inter-population phenotype distances (Section 2.4) and embed them as weights on every edge of a fully connected graph. When laid out using a force-directed algorithm (Figure 3A), populations with smaller distances naturally cluster closer together, but the sheer number of edges obscures the most informative biological relationships.

**Figure 3.**
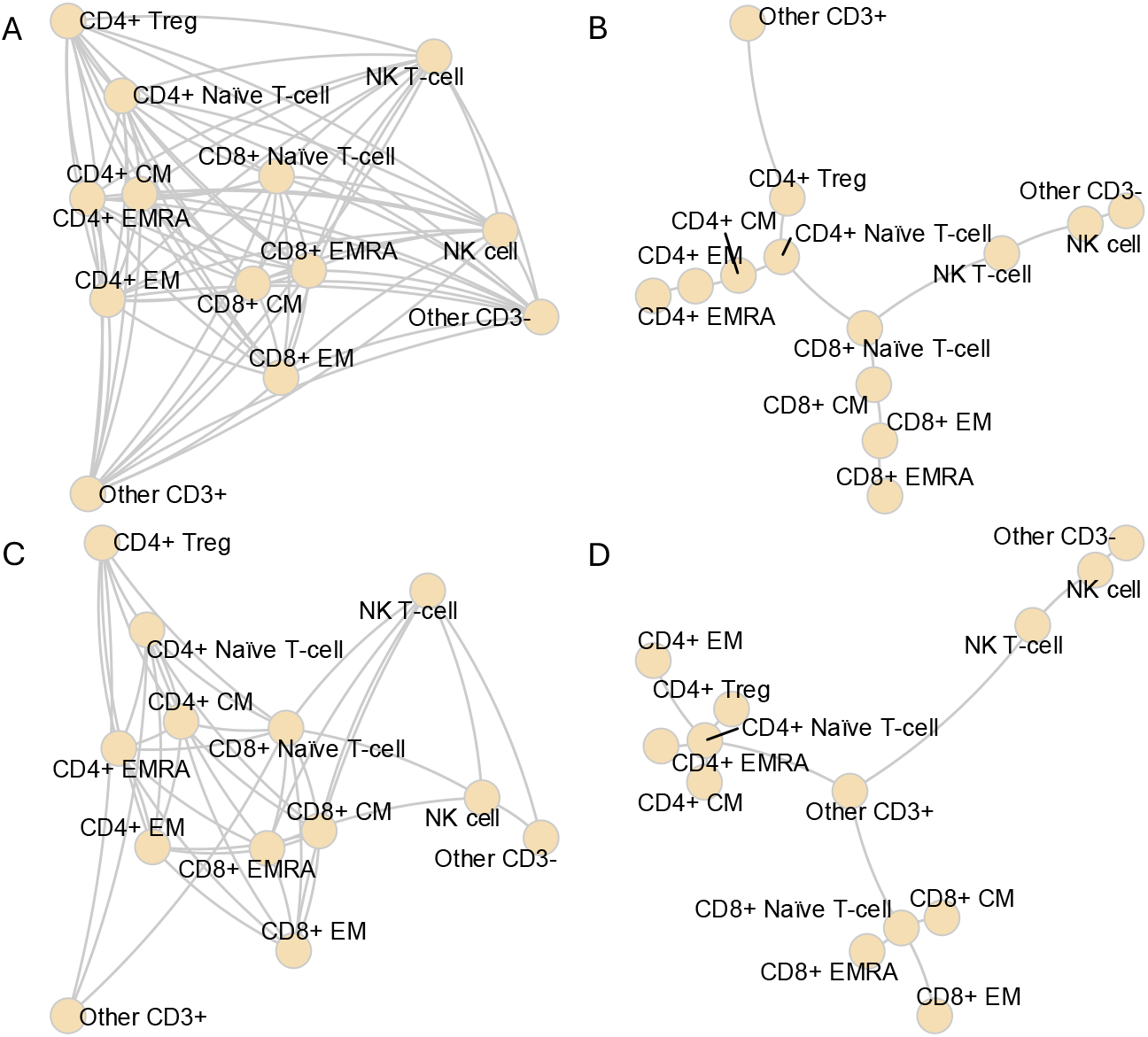
A collection of phenotype-aware layouts for the graph-based visualization of the NK cell and T-cell populations. (A) Fully-connected layout. (B) Minimum spanning tree layout. (C) Distance-threshold layout. (D)Expert-curated layout.

To distill the graph into a more interpretable structure, we explored three edge-filtering strategies. First, extracting a minimum spanning tree (Figure 3B) preserves the strongest links required to maintain connectivity without redundant cycles. Second, applying a distance threshold (Figure 3C) prunes all edges whose weights exceed a chosen cutoff (we use the average value of the edge weights for demonstration; other meaningful cutoffs can be chosen also), removing weaker associations. Third, expert immunological knowledge guides the selection of edges identifies as the most biologically meaningful (Figure 3D). For the remainder of this report, we adopt the expert-curated layout (Figure 3D) to ensure our visualizations emphasize the relationships most relevant to disease monitoring and treatment response.

### 3.1 Graph visualization and comparison of FC samples

The MPM dataset comprises flow-cytometry samples collected from each patient at three therapy milestones: baseline (pre-vaccination), two weeks after the first vaccine, and two weeks after the third vaccine. In Figure 4A, we visualize patient MCV005’s co-inhibition-panel samples at these three time points using our expert-curated graph layout. Here, each vertex represents one of the thirteen gated Lymphocyte populations, with its radius proportional to that population’s fraction of total lymphocytes. Vertex positions are determined by pairwise phenotype distances, so phenotypically similar populations lie closer together.

**Figure 4.**
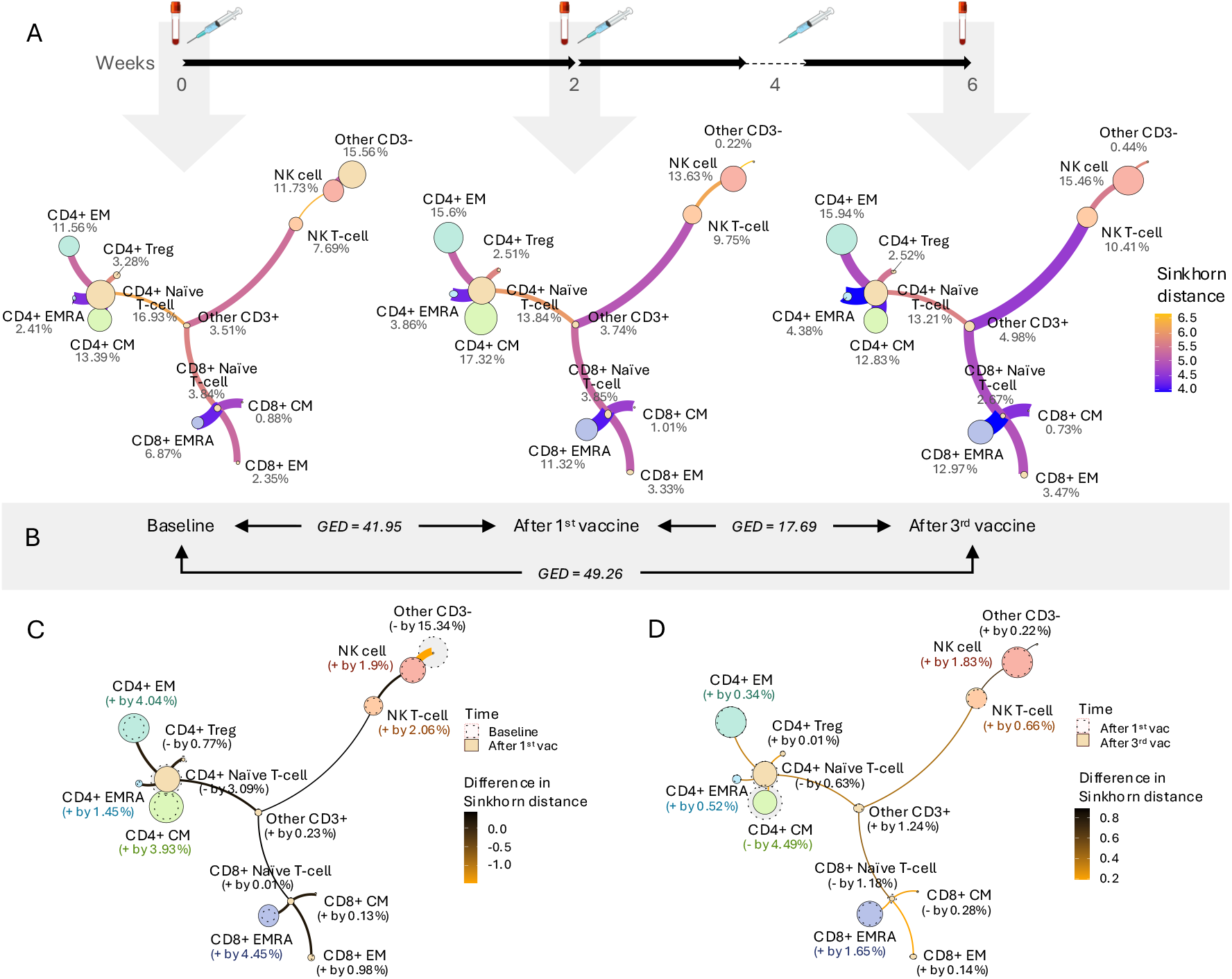
Sinkhorn distance-based visual summary of changes in T-cell populations in MPM patient due to dendritic cell-based immunotherapy (DCBI) (Patient ID: MCV005). (A) Illustrations of the flow cytometry sample graphs collected at baseline, two weeks after the first vaccine, and two weeks after the third vaccine using the edge-customized layout. (B) The graph edit distances (GED) between the sample graphs shown in A. (C-D) Visual comparison of sample graphs between baseline and after the first vaccine, and after the first and third vaccine.

Edges weight (but not length) encode the inter-population Sinkhorn distances: smaller distances (indicating greater similarity in marker expression) produce thicker, more vividly colored edges, while larger distances yield thinner, paler edges. Both vertex sizes and edge-weight scalings are normalized on a per-patient basis, ensuring that comparisons across the three time points reflect true biological shifts rather than arbitrary scaling differences.

It is worth noting that two populations may appear close together, reflecting phenotypic (functional) similarity or a lineage relationship, yet still differ in marker-abundance profiles, as indicated by a thinner connecting edge. Although Figure 4A displays individual FC samples, the same graph-based approach can be applied to averaged data from multiple samples (not shown). This yields a compact, informative visualization that combines biological context with statistical rigor.

From the three time-point graphs in Figure 4A, we observe trends consistent with Dietz et al. Dietz et al. (2023):

- Natural killer cells, effector memory (EM) T cells, and terminally differentiated effector memory (EMRA) T cells increase in proportion over the course of treatment.
- Naïve T cells decrease in abundance following the third vaccine.
- Central memory (CM) T cells rise after the first vaccine but decline after the third.

We also observe that several edges, for instance, the edge connecting NK cells and NK T-cells, get thicker and more vividly colored along the vaccination process. It indicates the inter-population Sinkhorn distance getting smaller, meaning the two populations are becoming more similar in their marker expression profiles. Biologically, this suggests that during the vaccination process, the marker expression patterns of these two populations, NK and NK T-cells, are converging, requiring less “effort” to transport one population into the other conceptually. This convergence may reflect shared or overlapping functional states emerging over time.

Although the sample graphs share the same layout and consistently scaled vertices and edges, comparing two graphs side by side can still be challenging, especially when changes in cell proportions or inter-populational Sinkhorn distances are subtle. To address this, we construct a comparison graph that overlays two FC samples using a shared layout, effectively highlighting their differences. Figures 4C and 4D present visual comparisons between the baseline and post-first vaccine samples, and between the post-first and post-third vaccine samples, respectively. In these comparison graphs:

- Vertices from the two samples are distinguished by style: one sample’s vertices appear as dotted circles and the other’s as solid circles. When a population’s proportion increases, the solid circle encloses the dotted one; when it decreases, the dotted circle is larger. Numeric labels at each vertex indicate the percentage change in that cell population.
- Edges encode the change in Sinkhorn distance between the two samples. A positive edge weight denotes increased similarity (i.e., a decrease in Sinkhorn distance), whereas a negative weight denotes decreased similarity (i.e., an increase in Sinkhorn distance).

This visualization style presents a succinct summary of the evolution of cell populations and their relationships between two time points, providing valuable insights into therapeutic effects. Furthermore, this approach extends beyond treatment monitoring, enabling comparisons between healthy and diseased samples or tracking disease progression, scenarios where specific cell populations may emerge or disappear entirely. Our graph-based methodology effectively accommodates such dynamic processes.

Beyond visual comparisons, we incorporate a quantitative measure of similarity between FC samples, known as graph edit distance (GED), as detailed in Section 2.5. Figure 4B presents the GED values for the sample graphs illustrated in Figure 4A. GED effectively captures changes in both cell proportions and inter-population Sinkhorn distances. While a single GED value may provide limited insight, a comprehensive analysis of GEDs across all three time points and patients within the three T-cell panels reveals more meaningful patterns, which will be further explored in Section 3.2.

Note that, while these graphs may visually resemble those generated by SPADE, the two approaches are fundamentally different in both methodology and purpose. SPADE clusters cells and builds a minimum spanning tree to infer cellular hierarchies, relying on density normalization and unsupervised clustering. In contrast, our graph-based visualization uses predefined cell populations and constructs phenotype-aware graphs, where edges are weighted by Sinkhorn distances to reflect distributional differences in marker expression. It also introduces graph edit distance (GED) for quantitative sample comparison, enabling robust tracking of biological changes, capabilities that SPADE does not natively support.

### 3.2 Relevance of graph-based visualization and graph edit distances for MPM dataset

In this section, we align our visualization and quantification framework with the key clinical observations of Dietz et al. (Dietz et al., 2023). Their trial evaluated adjuvant dendritic cell–based immunotherapy in 14 MPM patients following CRS–HIPEC, using six 14-color flow-cytometry panels sampled at baseline, two weeks post-first vaccine, and two weeks post-third vaccine. Although total lymphocyte and CD8^+^ T-cell frequencies remained stable, DCBI induced a robust memory T-cell response:

- **Effector memory (EM) and central memory (CM) T cells** expanded significantly after the first dose and remained elevated through the third.
- **Naïve T cells** declined sharply following the third vaccine.
- **CD8**^+^ **EMRA T cells** increased most notably in patients with prolonged progression-free survival.

Dietz et al. summarized these dynamics with grouped boxplots and stacked bar charts (see their Figure 2). While effective for trend analysis, these plots require multiple panels to cover all subpopulations and do not convey the phenotypic relationships among them.

Our graph-based representation (Figure 4) consolidates each sample into a single network:

- **Vertex size:** scales with each subpopulation’s proportion.
- **Vertex position:** determined by phenotype similarity (Hamming distance), so similar populations lie closer together.
- **Edge thickness and color:** encode Sinkhorn distances, capturing distributional shifts in marker expression.

This unified visualization not only reproduces the reported increases in EM and CM subsets, the decline in naïve cells, and the EMRA enrichment, but also reveals how these populations relate phenotypically, enhancing both statistical rigor and biological interpretability in high-parameter cytometry.

Importantly, cell subpopulations in a flow cytometry sample are not isolated - they are biologically interconnected. Simply tracking changes in individual cell percentages does not fully capture the complexity of the immune response. By incorporating changes in Sinkhorn distances between subpopulations, our graph-based method reflects underlying biological interactions. This provides a thorough understanding of immune modulation within and between samples.

We calculated graph-edit distances (GEDs) for each MPM patient at baseline, two weeks after the first vaccine, and two weeks after the third vaccine, uncovering an intriguing pattern (Figure 5A). In some patients, GEDs rose steadily from baseline to the third vaccination, while in others they fell after an initial post-first-dose increase. Because GEDs quantify shifts in cell-population structure, these trajectories suggest that certain patients mounted an enhanced immune response by the third vaccination, whereas others showed a diminished response.

**Figure 5.**
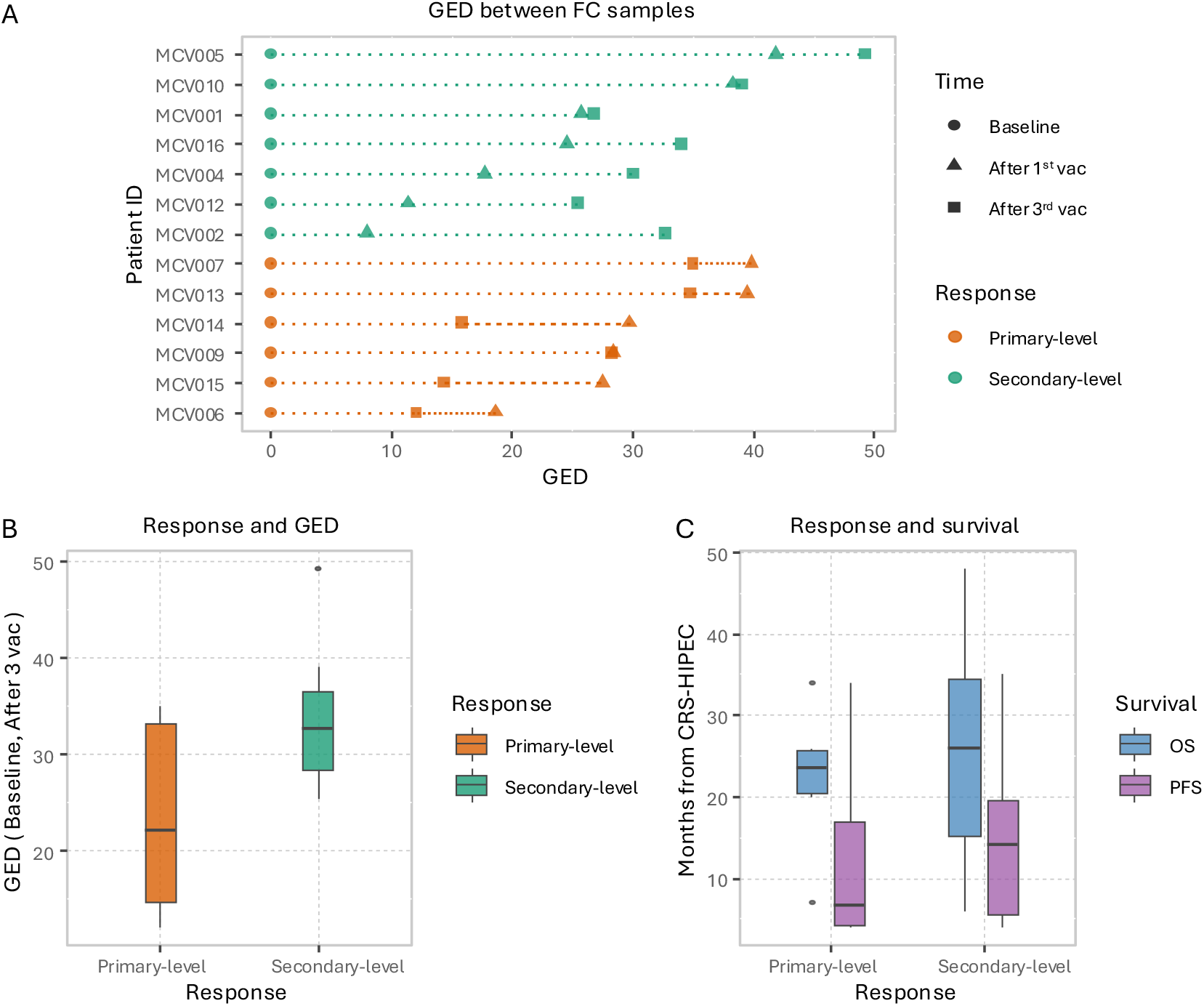
Graph edit distance (GED) insights and clinical outcomes in MPM patients. (A) GED trajectories across three vaccination time points for two distinct patient response groups. (B) Comparison of overall GED from baseline to the third vaccine between primary-level and secondary-level responders. (C) Comparison of progression-free survival (PFS) and overall survival (OS) between primary-level and secondary-level responders.

Based on these dynamics, we classified patients as primary-level responders whose GEDs declined following the third dose, or secondary-level responders whose GEDs continued to increase. Figure 5B shows that secondary-level responders exhibited a higher overall GED between baseline and the third vaccination than primary-level responders. When we compared these groups against clinical outcomes reported by Dietz et al. (2023), specifically progression-free survival (PFS) and overall survival (OS) after CRS-HIPEC, we found that secondary-level responders demonstrated longer PFS than primary-level responders (Figure 5C).

Due to high variability in the patients’ state at baseline, statistical significance of their immunological shifts by the immunotherapy regimen could not be verified. Therefore, we work under the assumption of normalized baseline GEDs across patients (Figure 5A) and performed a pseudo-F test on the changes in GED to the third vaccination. The result was statistically significant (*p* ≤ 0.001), confirming that the immunotherapy regimen likely produced substantial immunological shifts by the third dose.

## 4 Optimal transport-based visualization for AML dataset

To demonstrate our framework’s versatility beyond treatment monitoring in the MPM dataset, we applied it to flow cytometry data from a publicly available acute myeloid leukemia (AML) dataset to distinguish healthy from diseased profiles. This AML dataset comprises eight cytometry panels, each measuring five markers; here, we focus on panel 6 (HLA-DR, CD117, CD45, CD34, CD38) to identify AML blasts (myeloblasts), Monocytes, and Lymphocytes. Note that, the Lymphocytes and Monocytes gatings are confirmed by markers in panel 7. Given the smaller number of populations relative to the MPM study, we constructed a phenotype-aware minimum spanning tree (MST) for our graph-based visualization. Figure 6 presents MST-based graphs for a healthy donor (Individual 1), an AML patient (Individual 5), and their direct comparison. In the AML patient, lymphocyte proportions are markedly reduced, whereas the myeloblast population is prominently expanded. These findings align with the characteristic immunophenotype of AML.

**Figure 6.**
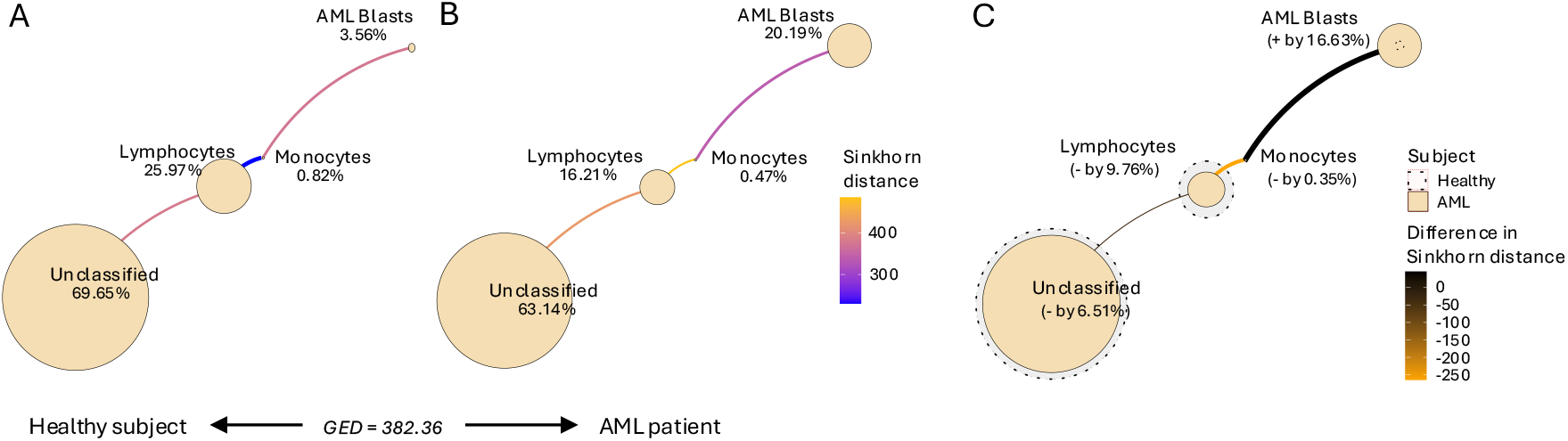
Sinkhorn distance-based visual summary of changes in cell populations in AML dataset using an MST-layout. (A) Illustration of an FC sample graph of a healthy subject (Individual 1). (B) Illustration of an FC sample graph of an AML patient (Individual 5). (C) Visual comparison of sample graphs between healthy and diseased. The graph edit distance (GED) between the graphs in A and B is shown.

## 5 Discussion

In the work described in this report, our primary objective was to develop interpretable, quantifiable visualization techniques for high-parameter flow cytometry. We deliberately de-prioritized the cell-classification step, working under the assumption that our visualizations would be applied downstream of diverse pre-processing pipelines. Accordingly, we used a straightforward, semi-automated gating procedure to assign cell populations, ensuring consistency with strategies previously applied to these data. While the implementation of advanced classification or clustering methods lies beyond our scope, we acknowledge that integrating our phenotype-aware layouts and inter-population Sinkhorn distances into existing frameworks (e.g., FlowSOM) could further enhance interoperability and analytical robustness of these flow-cytometry workflows.

From a computational standpoint, we employ the entropy-regularized Sinkhorn distance to quantify similarity (or dissimilarity) between cell subpopulations within each flow-cytometry sample. These pairwise distances are embedded as edge weights in a graph, enabling us to compare entire samples via graph-edit distance. Although one can directly apply OT to whole samples, bypassing the graph abstraction, such an approach poses several major challenges. First, solving a single OT problem between high-parameter samples is both computationally intensive and memory-hungry. Second, integrating cell-population and phenotypic information into the transport cost requires a careful, bespoke formulation. Third, the interpretability of OT solutions is severely limited when specific functional populations are not explicitly identified, necessitating a dedicated meta-layer framework for automated interpretation and translation of global OT results into specific population-level changes.

We acknowledge these obstacles and plan to explore a structured, end-to-end OT framework for sample-to-sample comparison in future work. Additionally, we intend to investigate alternative inter-population similarity measures, such as Marker Enrichment Modeling (MEM) (Diggins et al., 2018), which quantitatively characterizes cell populations based on marker enrichment relative to a reference. Integrating MEM-derived enrichment scores into our graph structures may offer a more interpretable and biologically grounded approach to analyzing graph edit distances, thereby improving our understanding of population-level changes across samples.

## 6 Conclusions

We have introduced a novel, biologically grounded framework for visualizing high-parameter flow cytometry data by utilizing OT theory. Through the use of the Sinkhorn distance, we quantified inter-population similarities and encoded these relationships into a graph-based visualization. Our approach combines phenotype-aware layouts with Sinkhorn distances to deliver compact, quantitatively rigorous summaries of cell-population structure, allowing direct comparisons across samples.

When applied to a dendritic cell–based immunotherapy trial in malignant peritoneal mesothelioma, our framework faithfully reproduced previously recognized immunological trends and alterations (e.g., memory T-cell expansion and naïve T-cell decline) and uncovered other potentially informative patterns of patient stratification. In particular, the strong association between GED results and progression-free survival underscores our method’s potential as a tool for immune monitoring and therapeutic decision-making in longitudinal studies.

Overall, this work bridges a mathematically rigorous approach and biological interpretability, laying the groundwork for actionable insights in clinical cytometry and immunology research.

## Appendix

The following material is from our previous work on the optimal transport framework (Shemonti et al., 2023).

### Optimal transport framework

The *transport problem* distributes a certain amount of *mass* from a set of sources to a set of destinations at minimum cost. There are two major factors in a transport problem: the *cost function* and the *transportation plan*. The cost function defines a fixed, non-negative effort required to transport unit mass from a source to a destination. This cost may only depend on the distance between the source and the destination or on other additional factors; in the former case, a Euclidean distance matrix between the sources and the destinations is a reasonable representation of effort.

Once the cost of transportation is represented, the remaining part of the problem involves transporting a non-negative amount of mass between sources and destinations, as described by a transportation plan. Various transportation plans result in different total costs, and the *optimal transport (OT) problem* aims to minimize this cost. The OT problem is *balanced* if the total mass at the sources equals the total mass at the destinations, and *unbalanced* otherwise (Peyré and Cuturi, 2019).

Let *r* and *c* be two *d* dimensional vectors representing the amount of mass at the sources and the destinations, respectively. The number of sources and destinations could differ, but they can be considered equal without loss of generality. Let *U* (*r, c*) be the set of all non-negative *d×d* matrices with row and column summing to *r* and *c*, respectively. Any matrix *P ∈ U* (*r, c*) describes a transportation plan that transports the *P* is ∑_*i,j*_ *P*_*ij*_*M*_*ij*_. Thus the OT problem between *r* and *c* given cost *M* can be formulated by Equation 1, where *D*_*M*_ (*r, c*) is the optimal transport distance:

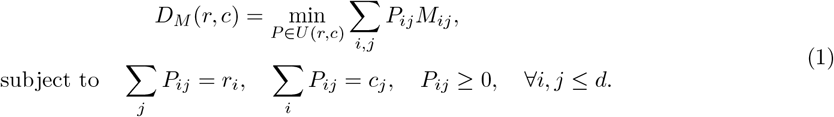

The masses in *r* and *c* could be normalized to sum to one, and then both *r* and *c* can be interpreted as probability distributions.

For *D*_*M*_ (*r, c*) to be a metric, the cost matrix *M* has to be a metric matrix (Villani, 2009; Avis, 1980; Brickell et al., 2008) satisfying the conditions shown in Equation 2.

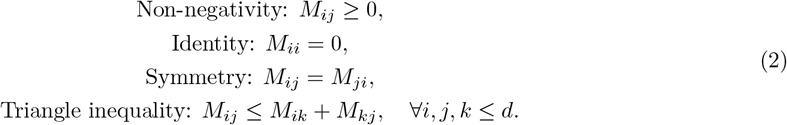

The OT is a convex optimization problem that can be solved using various approaches (Ahuja et al., 1993; Orlin, 1993). For a general cost matrix, the computational cost scales as 𝒪 (*d*^3^log(*d*)) (Pele and Werman, 2009), which prevents scaling the solution to large problem sizes. Earlier approximate solutions obtained by putting constraints on the cost matrix could result in a loss of applicability and performance (Grauman and Darrell, 2004). A later approximation to the original OT problem using an entropic regularization scheme was proposed by Cuturi (2013) to reduce the computational complexity. The scheme employs the Sinkhorn-Knopp matrix scaling algorithm (Sinkhorn and Knopp, 1967; Knight, 2008), and hence the name *Sinkhorn distance* for its objective function.

### Sinkhorn distance

A straightforward way of thinking about a transportation plan is by noticing that if a source contains more mass, it should originate more, and if a destination requires more mass, it should receive proportionally more. Such a transportation plan is represented by *rc*^*T*^, and the optimal plan *P* should be somewhere around the distribution *rc*^*T*^. Simply speaking, the idea of the entropic regularization scheme by Cuturi (2013) is to choose *P* from a smaller set near *rc*^*T*^, instead of the entire set *U* (*r, c*).

To capture these ideas, Cuturi (2013) imposes an additional constraint of Kullback-Leibler (KL) divergence on the OT formulation, as shown in Equation 3, and computes the Sinkhorn distance 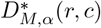. This constraint introduces a set *U*_*α*_(*r, c*) ⊂ *U* (*r, c*) from which an optimal transportation plan *P* is selected. The KL divergence distance between *P* and *rc*^*T*^ is set to be smaller than a predefined parameter *α*. In other words, *P* should belong to a distribution near *rc*^*T*^.

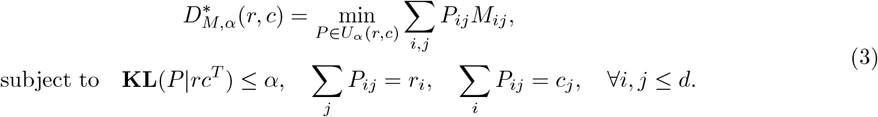

The entropy (*h*) of the transportation plan (*P*) and the mass vectors (*r* and *c*) are given in Equation 4:

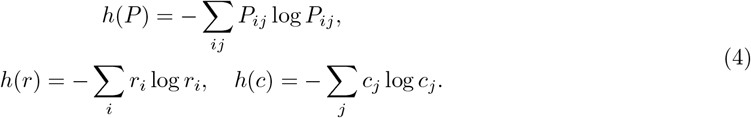

We proceed to express the KL divergence constraint in terms of the entropy:

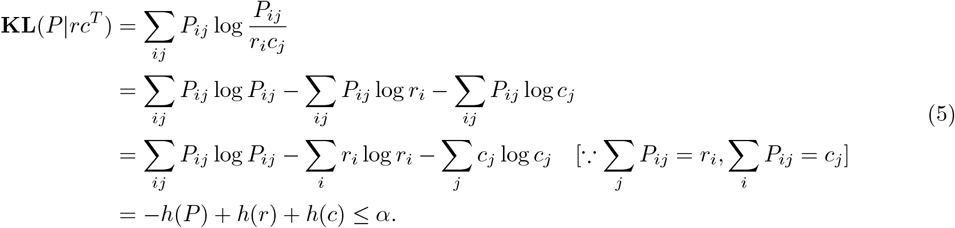

Thus, the new constraint states that the entropy of *P* should be large enough to satisfy

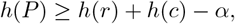

which constrains *P* to be chosen from the Kullback-Leibler ball of level *α* centered about *rc*^*T*^ (see Figure 1 in (Cuturi, 2013)).

This interpretation makes the OT problem non-convex, and an alternative formulation of Sinkhorn distance is required for ease of optimization. For every pair (*r, c*), each *α* corresponds to a Lagrange multiplier *λ ∈* [0, ∞) such that 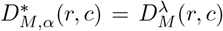 The distance 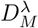, shown in Equation 6, is called the dual-Sinkhorn divergence by Cuturi (2013).

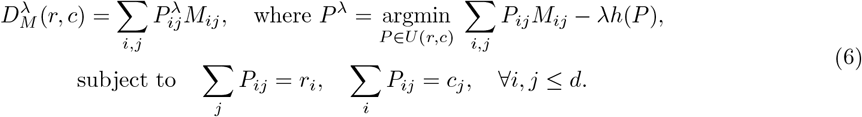

By introducing two dual variables *ϕ* and *ψ* for each of the two equality constraints of Equation 6, the Lagrangian of the objective function can be written as Equation 7.

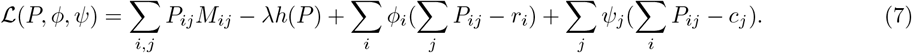

The derivative of the Lagrangian objective function with respect to *P*_*ij*_, for any pair (*i, j*), can be set to zero to obtain an extremum; the second derivative of the Lagrangian, 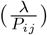, is positive since both the numerator and the denominator are positive, and thus we have obtained a minimizer of the Lagrangian.

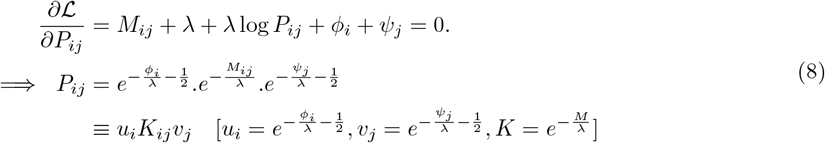

Given *K, r* and *c*, the Sinkhorn-Knopp matrix scaling algorithm converges to a solution *P* ^*λ*^ of the following form:

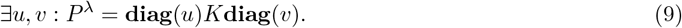

*P* ^*λ*^ should have the correct row and column sums, as shown in Equation 6. We deduce the update rule for the Sinkhorn-Knopp algorithms from those constraints in the following manner:

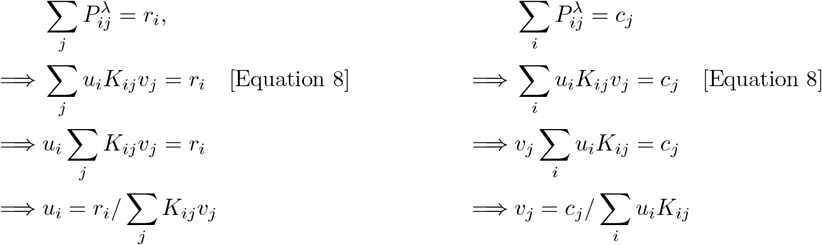

Thus the update rule for the Sinkhorn-Knopp algorithm can be written as Equation 10, where *v* can be initialized randomly.

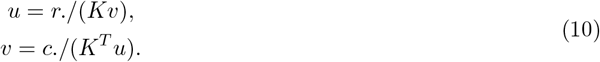

Cuturi (2013) observes that the number of iterations in the Sinkhorn-Knopp algorithm is bounded independent of *d*. Thus, the cost of computing 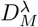 is *O*(*d*^2^), which is an improvement over *O*(*d*^3^log(*d*)). Cuturi (2013) describes an approach to compute the Sinkhorn distance 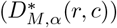 through the dual-Sinkhorn divergence 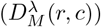, and also reports that the dual-Sinkhorn divergence does not perform worse than the classic optimal transport distances. Therefore, we use the dual-Sinkhorn divergence to measure the distance between the spatial statistics of the point patterns in our experiments and refer to as the Sinkhorn distance. We utilize the *POT Python Optimal Transport* library for computing the dual-Sinkhorn divergences (Flamary et al., 2021).

